# Evaluation of BMP-mediated patterning in a 3D mathematical model of the zebrafish blastula embryo

**DOI:** 10.1101/585471

**Authors:** Linlin Li, Xu Wang, Mary C. Mullins, David M. Umulis

## Abstract

Bone Morphogenetic Proteins (BMPs) play an important role in dorsal-ventral (DV) patterning of the early zebrafish embryo. BMP signaling is regulated by a network of extracellular and intracellular factors that impact the range and signaling of BMP ligands. Recent advances in understanding the mechanism of pattern formation support a source-sink mechanism, however it is not clear how the source-sink mechanism shapes patterns in 3D, nor how sensitive the pattern is to biophysical rates and boundary conditions along both the anteroposterior (AP) and DV axes of the embryo. We propose a new three-dimensional growing Partial Differential Equation (PDE)-based model to simulate the BMP patterning process during the blastula stage. This model provides a starting point to elucidate how different mechanisms and components work together in 3D to create and maintain the BMP gradient in the embryo. We also show how the 3D model fits the BMP signaling gradient data at multiple time points along both axes. Furthermore, sensitivity analysis of the model suggests that the spatiotemporal patterns of Chordin and BMP ligand gene expression are dominant drivers of shape in 3D and more work is needed to quantify the spatiotemporal profiles of gene and protein expression to further refine the models.

## 1. Introduction

Pattern formation by morphogens drives the normal development of various processes such as limb development and organogenesis in animals (Maini and Solursh 1991; Lecuit and Cohen 1998; Sansom and Livesey 2009; Rogers and Schier 2011; Rushlow and Shvartsman 2012; Bökel and Brand 2013; Tuazon and Mullins 2015). In zebrafish, patterns of gene expression along the dorsal-ventral (DV) body axis are regulated by Bone Morphogenetic Proteins (BMPs) (Tucker et al. 2008). BMPs are a member of the TGF-*β* (transforming growth factor *β*) superfamily. Very early in embryonic development, both invertebrates and vertebrates require BMP signaling to pattern the DV axis (De Robertis and Sasai 1996; Holley and Ferguson 1997). BMPs pattern DV tissues of zebrafish, *Xenopus*, and *Drosophila* embryos by forming a spatially-varying distribution, in which different levels of BMP signaling drive differential gene expression (Little and Mullins 2006).

BMP signaling is propagated by the binding of BMP dimers to serine/threonine kinase receptors on the cell membrane. Type I and II receptors form higher order tetrameric complexes and phosphorylate intracellular Smads (P-Smad5 in zebrafish) that accumulate in the nucleus and regulate differential gene expression. BMP signaling is regulated by different molecules at multiple levels: extracellular, intracellular, and on the membrane (Wang et al. 2014). These regulators form a system that enhances, lessens or refines the level of BMP signaling. One group of regulators are the inhibitors of BMP signaling (Little and Mullins 2006; Umulis et al. 2009; Tuazon and Mullins 2015). BMP inhibitors include Chordin (Chd), Noggin (Nog), Crossveinless2, Follistatin, Sizzled, and Twisted gastrulation. (Khokha et al. 2005; Dal-Pra et al. 2006; Umulis et al. 2009; Wagner et al. 2010; Dutko and Mullins 2011), most of which act by binding BMP ligands, preventing them from binding their receptors. In this study, we focus on the major antagonists Chordin and Noggin. Chordin, unlike Noggin and Follistatin, can be cleaved by the metalloprotease Tolloid, releasing Chordin-bound BMP ligand and allowing it to bind receptors and signal (Blader et al. 1997; Piccolo et al. 1997). Downstream intracellular regulation of BMP signaling occurs throughout the BMP-Smad pathway (von Bubnoff and Cho 2001). Inhibitory Smads modulate BMP signaling, either by interacting with Type I BMP receptors or by preventing R-Smads from binding Smad4. Others molecules, such as microRNAs and phosphatases may also act as intracellular modulators (Luo et al. 2015).

In our previous work, we developed a data-based 1-D model to investigate the mechanisms of BMP-mediated DV patterning in blastula embryos to early gastrula embryos at 5.7 hours post fertilization (hpf) before the initiation of BMP-mediated feedback (Zinski et al. 2017). We quantified BMP signaling in wild-type (WT), *chordin* mutant, and *chordin* heterozygous embryos. In our previous model screen, we simulated BMP gradient formation along a 1D line at the embryo margin and used the quantitative measurements of P-Smad to inform our model selection. We concluded that the signaling gradient patterning the vertebrate DV axis is most consistent with either a counter gradient or a source-sink mechanism. Measurements of BMP2 diffusion by fluorescent-recovery after photobleaching of ∼4um^2^/sec by ourselves and others, in addition to published estimates for the BMP2 lifetime, were more consistent with the source-sink mechanism than the counter gradient mechanism (Pomreinke et al. 2017; Zinski et al. 2017).

These models lay the ground work for our current study where we seek to add growth and 3D patterning to the model and test mechanisms of patterning in 3D. There is a significant need for spatially and temporally accurate 3D models of the early embryo to evaluate reaction-diffusion processes of chemical morphogens including BMP ligands. The development of accurate 3D models has been limited due to the complexity of embryo structure, and the computational resources needed to run parametric screens in 3D models. The work here builds off of previous work carried out by us and others. Previous work in Drosophila focused on a three-dimensional organism-scale model of BMP and gap gene patterning in the *Drosophila* embryo (Umulis et al. 2010; Hengenius et al. 2011; Umulis and Othmer 2015). In zebrafish, the 3D structure shares some similarities but many differences with *Drosophila*, especially in regard to the growth during epiboly and the potential role of cell movement in shaping the gradient in the zebrafish. To date there are few mathematical models for zebrafish embryonic development and only one that developed a 3D approximation by Zhang et al. to study the role of Chordin in regulating BMP signaling in zebrafish (Zhang et al. 2007). This model focused on the blastula stage from 30% epiboly to the early gastrula shield stage (around 50% epiboly), however, it did not include growth, was not compared to data, and suggested a mechanism of BMP shuttling that has now been shown not to function in the embryo (Zinski et al. 2017).

BMP signaling begins patterning ventral tissues before the onset of gastrulation (Tucker et al. 2008; Bhat et al. 2013; Hashiguchi and Mullins 2013). During gastrulation, coordinated cell movements organize the germ layers and establish the primary body axes of the embryo (Lepage and Bruce 2010). Epiboly begins during mid blastula stages (30% - 50% epiboly) and continues through the gastrula stage and entails a thinning and expanding of the cell layers from the animal pole, where the blastoderm lies, to cover the yolk cell over the vegetal pole (Warga and Kimmel 1990). The 3D model developed here accounts for this growth from 30% to 50% epiboly. The cell proliferation and movement during epiboly leads to a growth-like cell flow in the system. Such systems are often considered on a growing domain and many have incorporated domain growth into models of pattern formation (Crampin et al. 1999; Klika and Gaffney 2017). There are many ways and levels of detail that can be developed into a “growing” domain in a model and the appropriate choice depends on the physics of growth, the relative rates for new mass to enter the system, the ratio of processes that are involved in transport of the molecules of interest among other considerations (Madzvamuse et al. 2005; Fried and Iber 2014). In our first model for early zebrafish development that encompasses up to 50% epiboly, the approaches we considered include finite element modeling with a moving mesh or a simplified approach that simply adds mass at the leading edge to represent the movement of the cells further in the vegetal direction using a finite-difference solution approach. In this study, we used a growth domain finite-difference approach that treat the growth of the 3D simulation domain by adding new layers of mass near the leading edge of cells at the margin of the embryo at each growth step and exclude advective flow since the system through 50% epiboly is likely diffusion dominant.

While our final goal is to develop a complete advection-diffusion-reaction model that incorporates all stages of zebrafish embryonic development, our current data and the development of the model herein that is tested against the data covers the first half of epiboly up through 50% epiboly. It also serves as a greater test of the source-sink mechanism. Herein, we find that a Partial Differential Equation (PDE)-based model for BMP patterning of a zebrafish embryo with growth through epiboly demonstrates that the source-sink mechanism patterns well in 3D, however, sensitivity analysis suggests that the prepatterns of mRNA expression of BMP ligands and the inhibitor Chordin play a large role in dictating the overall shape and dynamics of the BMP gradient. Additional work is needed to quantify the gene expression domains and map them into the 3D modeling environment to improve the model for greater understanding of the inhibitors’ roles in shaping the gradient in 3D.

## 2. Method

We adopted our previous 1D model for the reaction-diffusion system simulated along a line for the margin (Zinski et al. 2017) to our 3D geometry for the embryo growing domain model. The regulatory network in this study includes four individual components (BMP, Chd, Nog, Tld) and two bound complexes (BMP-Chd (BC), BMP-Nog (BN)). Figure 1H illustrates the regulatory network between different components: antagonists Chd and Nog inhibit BMP signaling by binding BMP ligands, and Chordin and BC complexes are cleaved by the metalloprotease Tolloid. To develop the model equations, a number of assumptions and simplifications are needed that are based on the biology and the questions being investigated. At minimum, we needed to determine if the mass-balance equations require both advective and diffusive terms for molecular transport, the structure of the reaction rate equations, and quantification of the sources of each of the components that may change over space and time.

**Figure 1.**
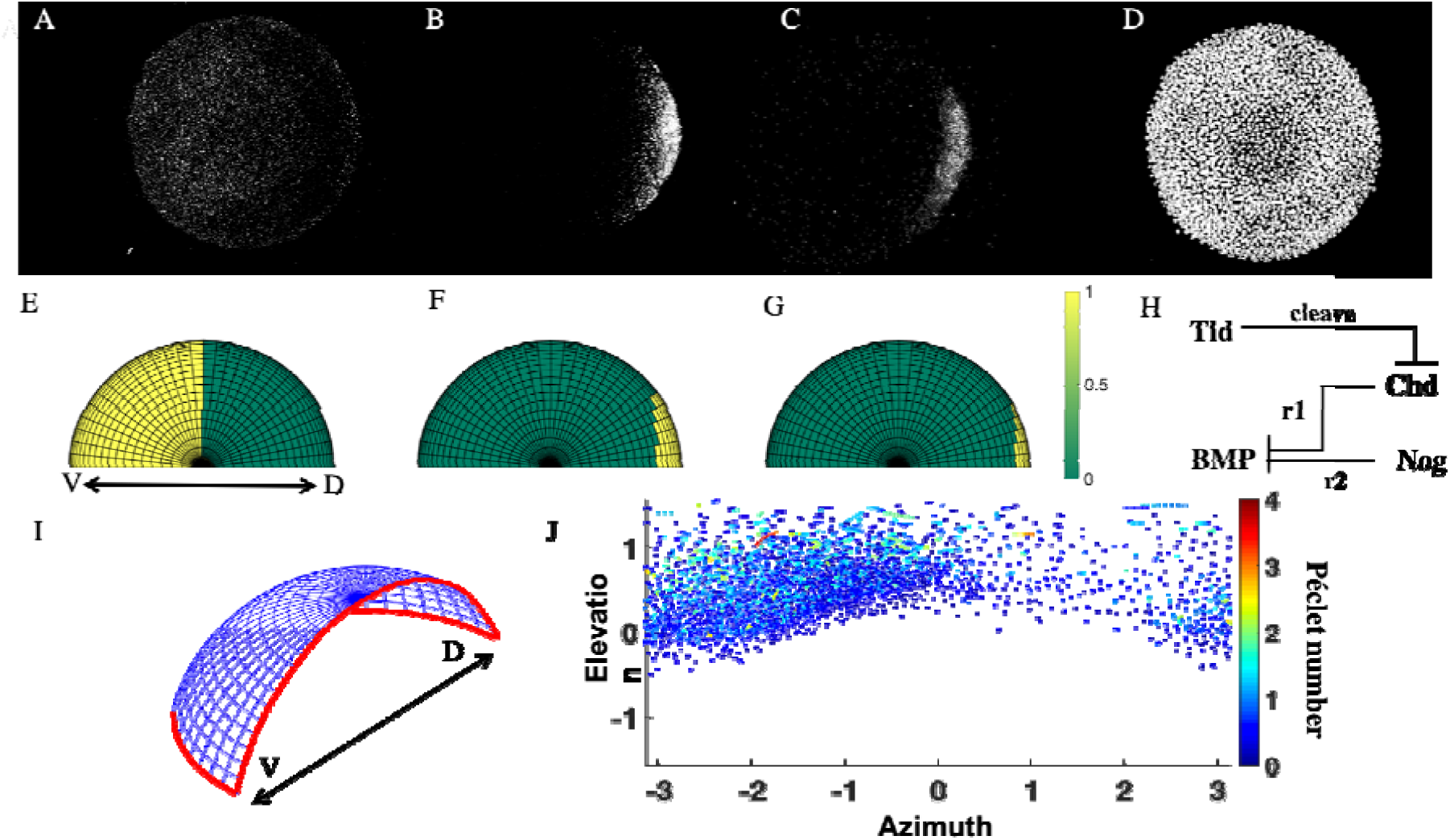
Whole mount experimental result of *bmp2b* (A), *chd* (B), *nog* (C) mRNA expression and nucleus positions (D), view from the animal pole. (E-G) Simulation input of expression area in WT for BMP, Chd and Nog, respectively. Yellow color indicates an area of production for each species. (H) Simplified BMP regulatory network used in this study. (I) Spatial domain at 3.5 hpf, Red open edges represents the non-flux boundary condition. (J) Cell trace map on elevation and azimuth direction around 50% epiboly (4.7-5.3 hpf), color represents the Péclet number based on the cell velocity calculated every 15 min.

We first estimated the potential role of advection in shaping the gradient based on our estimates for diffusion and the rates of cell movement in early development. We quantified the cell movement during epiboly by using the digital embryo data presented by Keller et al. (Keller et al. 2008) and quantified individual cell traces from 30% to 50% epiboly. To decide whether the advective transport caused by the cell movement or diffusive transport dominates the BMP concentration profile during blastula stages, we estimated the average Péclet number based on the cell tracing data from 4.7 to 5.3 hpf.

The Péclet number (Pe) is a dimensionless number that represents the ratio of the convection rate over the diffusion rate in a convection-diffusion transport system (R. Byron Bird Warren E. Stewart Edwin N. Lightfoo et al. 2006). The Péclet number (in terms of time-scales for each process) is defined as:

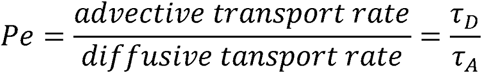

where 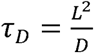 and 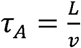, Thus

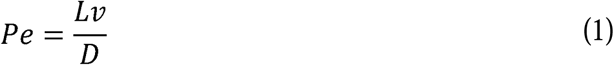

*v* represents linear flow velocity; L represents the characteristic length of the flow and D is the diffusion constant. In this study, one-quarter of the embryo radius was used as the characteristic length due to the average distance traveled for components considering the spatially distributed sources. Figure 1J illustrates the cell trace by 5.7 hpf on a 2D map of elevation and azimuth directions, the color scale represents the Péclet number based on the cell velocity. The median level of Péclet number among all the trackable traces is 0.428 during the blastula stage, and close to the margin (where the patterning considers happening), it is even smaller (median level 0.38), supporting diffusion dominant transport prior to 50% epiboly. Later during gastrulation, we found that the Péclet number is approximately or larger than 1 throughout the entire embryo, suggesting a need for simultaneous advection and diffusion. With a Péclet number in the measured rates for blastula stage, the time scale for diffusion is about 2-3 times lower than for advection, suggesting that the advective term is a minor contributor to flux. Thus, for the blastula stage embryo, we can assume this problem as a moving domain non-advection problem.

Based on the regulatory network in our model (Figure1H), we developed five-coupled non-linear partial differential equations (PDEs) for BMP ligand, Chordin, Noggin, and the complexes of BMP-Chordin, BMP-Noggin. The model solves the spatial-temporal diffusion problem in spherical coordinates using the derivative in expanded form:

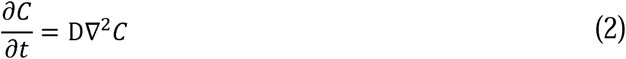

into:

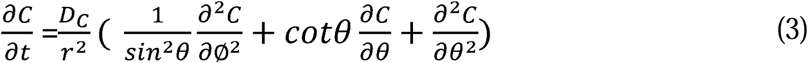

Note, C here represents a generic concentration, where C in the following equations represents the concentration of Chordin. The reaction between the different components are list below:

The reaction of BMP ligand and Chordin ligand forming BMP-Chordin (BC) complex,

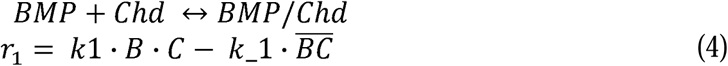

BMP ligand and Noggin form a BMP-Noggin (BN) complex,

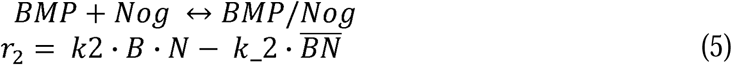

the metalloproteases Tolloid (Tld), which cleave and inactivate Chd, and BMP-Chd complex. *k*_*n*_ and *k*_−*n*_ are the forward and reverse reaction rates, respectively.

The model for all the species involved is given by **Equations (6-10 below)**

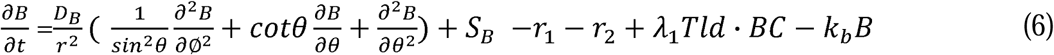

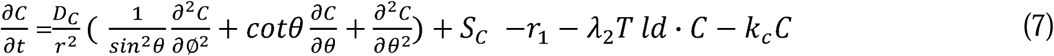

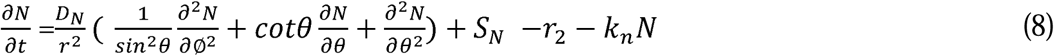

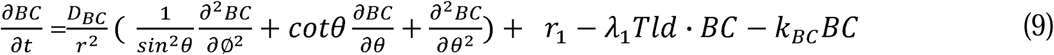

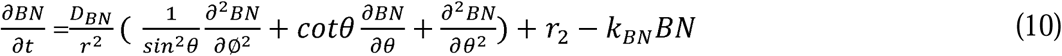

Since we limited the domain to the sphere surface, the diffusion in the r direction can be ignored. The above equation holds in the growing spatial domain *Ω*_*t*_ for all 0 < *t* < *T*, where *ϕ* and θ ∈ *Ω*_*t*_. B, C, N, BC, BN and Tld represent the concentration of BMP, Chordin, Noggin, BMP-Chordin complex, BMP-Noggin complex, and Tolloid, respectively. *D*_*i*_ represents the diffusion rate for each species, *S*_*i*_ is the constant term of source for original expression of the specific gene which varies by its spatial distribution and reflects the experimental gene expression data for different species (Figure 1E, 1F, 1G), *λ*_*i*_ is the Tld processing rate for Chd and BC complex, and *K*_*i*_ is the decay rate for specific ligands.

### Measurement of source distributions

We use the range of mRNA expression to represent the protein secretion of different species. To determine the values for source terms *S*_*B*_, *S*_*C*_, and *S*_*N*_ in the model, we imaged the spatial domains for expression of *bmp, chd*, and *nog* mRNA throughout the embryo using the RNAScope method. The embryos were fixed at the desired developmental stages with 4% PFA at RT for 4 hours or at 4°C for 24 hours and washed with 0.1% PBSTween 3 times at RT, each for 10 min. 20-30 embryos were separated into 1.5ml RNase free tube. The chorions were removed, and the embryos were gradually dehydrated from 25% methanol in PBST, 50% methanol in PBST, 75% methanol in PBST to 100% methanol, each for 5 min at RT. Store the embryos at −20 °C at least one day and up to 15 days. Two drops of Pretreat 3 (ACD, #320045) were added at RT for 15 min to permeate the embryos. Counterstaining of the probes using the RNAscope Fluorescent multiplex detection reagents (ACD, #320851). The user’s manual (323100-USM) is available online, however, we made some modifications. We performed probe hybridization at 40 °C for 16 hours, and C2 probes need to be diluted by C1 probes. Detailed information about the probes is shown in Table 1. All wash steps were performed three times using 0.2x SSCT for ten minutes each time. DAPI was used to stain the nuclei at 4 °C overnight. We chose AltC for Amp4 in the staining kit.

**Table 1.**
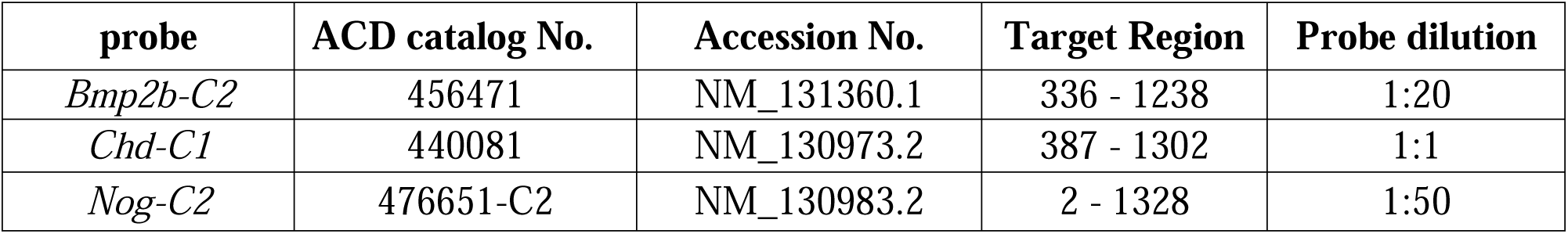
RNA Probe Information.

Embryos were mounted in 1% low melting agarose on 35mm glass bottom microwell dishes (Matek, P35G-1.5-10-C). Whole-mounted embryos were imaged with a 20×/1.0 Plan-Apochromat water immersion lens (D = 0.17 M27 75 mm). *chd* mRNA was imaged by excitation at 555 nm wavelength. *bmp2b* and *nog* mRNA were imaged by excitation at 647 nm wavelength.

We approximated simple expression regions from confocal images to build the model. Various expression regions were tested for all three components (BMP, Chd, and Nog). Figure 1 illustrates a set of expression regions for BMP, Chd, and Nog that were tested. For the simulations, the image data were converted into binary regions of expression and no expression. As shown in Figure 1, the BMP production region is limited to the ventral region, and Chd and Nog are limited to the dorsal side. As shown in Figure 1C, the Nog production region is smaller than the Chd production region.

The zebrafish embryo is approximated as a perfect hemisphere and the reaction-diffusion process happens on the surface of the sphere. Solutions were computed using the finite difference scheme based on the elevation and azimuth angles in spherical coordinates. Since the experimental result indicates that the development of zebrafish embryo is symmetrical during the blastula and gastrula stage, we only calculate our model in a quarter-sphere domain to decrease the computation and storage load. No-flux boundary conditions were applied for all species on both the ventral and dorsal boundaries (Figure 1I). To avoid the singularity that happens on the top points where the elevation angle *θ* is equal to zero, we used a hollow-shaft approximation method presented by Thibault et al (Thibault et al. 1987). In which mesh point *θ* = 0 is eliminated by introducing a small but finite interior surface *θ* = *f*_*θ*_ Δ*θ*, we have tested and used *f*_*θ*_ = 0.1 in this study. The growth of the domain is achieved by adding finite-difference layers at the marginal region on elevation direction. Mass is added to the system by matching the concentration of the newly added node with the closest margin node of the previous time point. Example geometries and a time-lapse of a single solution are shown in Figure 2A.

**Figure 2.**
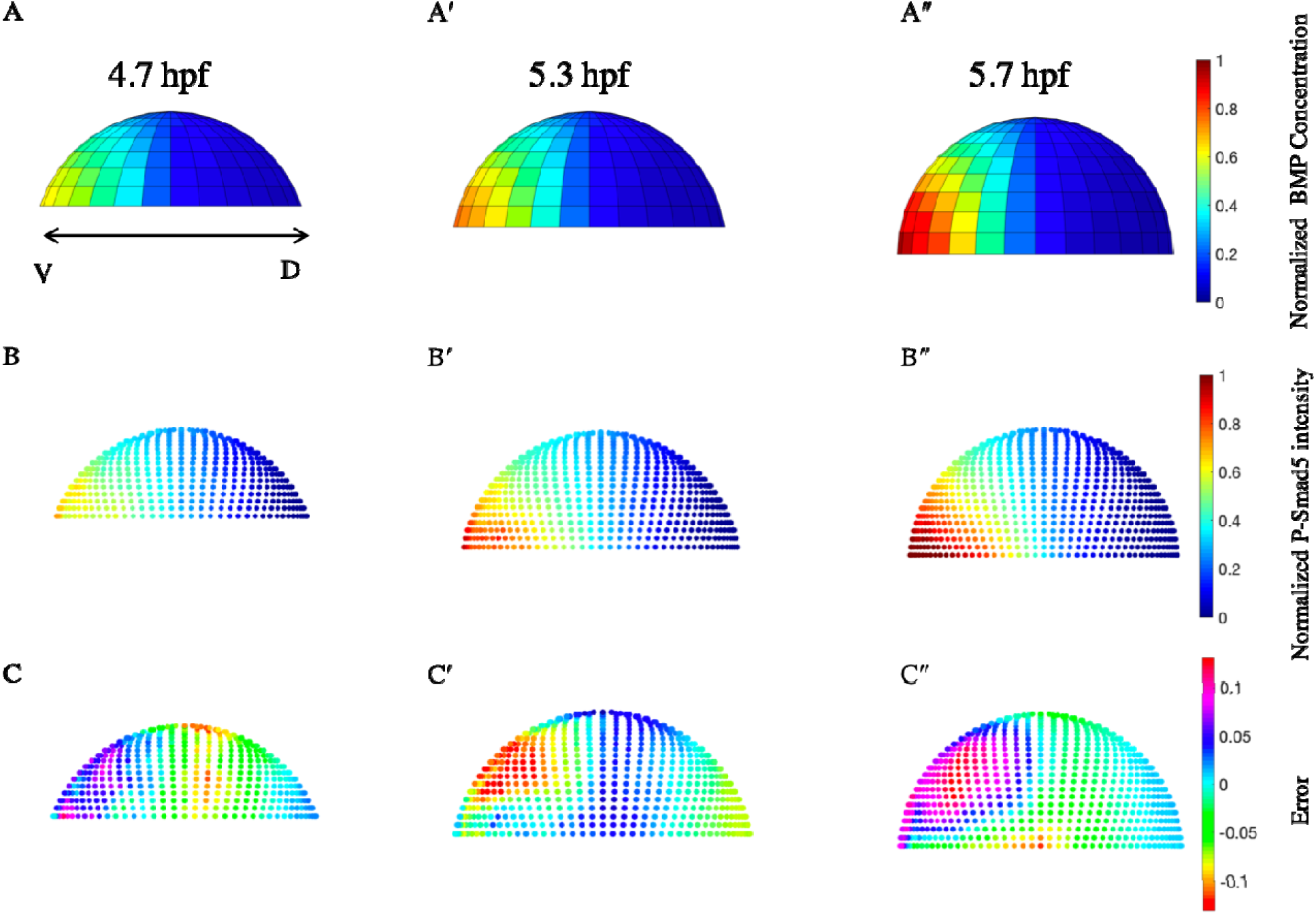
A, A’ and A”, Normalized simulation result of a wild type case at 4.7 hpf, 5.3 hpf, and 5.7 hpf. B, B’ and B”, Average and normalized P-Smad5 profile of 4.7 hpf, 5.3 hpf, and 5.7 hpf embryos. C,C’ and C”, Relative differences between simulation results and P-Smad5 level. Positive error indicates the experimental data are higher than simulation results, negtive error indicates the experimental data are lower than simulation results.

To test the Partial Differential Equation (PDE)-based models developed herein, we used the previously published point cloud data for P-Smad5 (readout of BMP signaling) that is available online at Zinski et al. (Zinski et al. 2017). To apply these data to our model, we processed the data by fitting the original data to a standard size hemisphere with different levels of coverage along the elevation direction based on the embryonic stage of development. Different sets of experimental data in the same stage were averaged on the sample points over the globular domain to obtain the representative P-Smad level for each stage as shown in Figure 2B.

To evaluate the system over a wide range of possibilities, we developed a computational model-based screen of over 300,000 combinations of biophysical parameters of the major extracellular BMP modulators. The parameter sets were randomly sampled in the parameter space listed in Table 1. Each parameter combination was then re-simulated without Chordin to predict the BMP signaling gradient in a *chordin* loss-of-function (LOF) scenario. Based on previous studies (Pomreinke et al. 2017; Zinski et al. 2017), the diffusion rate and decay of BMP ligand and Chd are fixed as constant, and for BMP, and and for Chd, respectively. The parameter space for the rest of unknown parameters consist of the range used in Zinski et al’s study (see Table 2). The model is solved for the developmental window that spans from 3.5 to 5.7 hpf, and all measurements of model error are calculated at 4.7, 5.3, and 5.7 hpf.

**Table 2.**
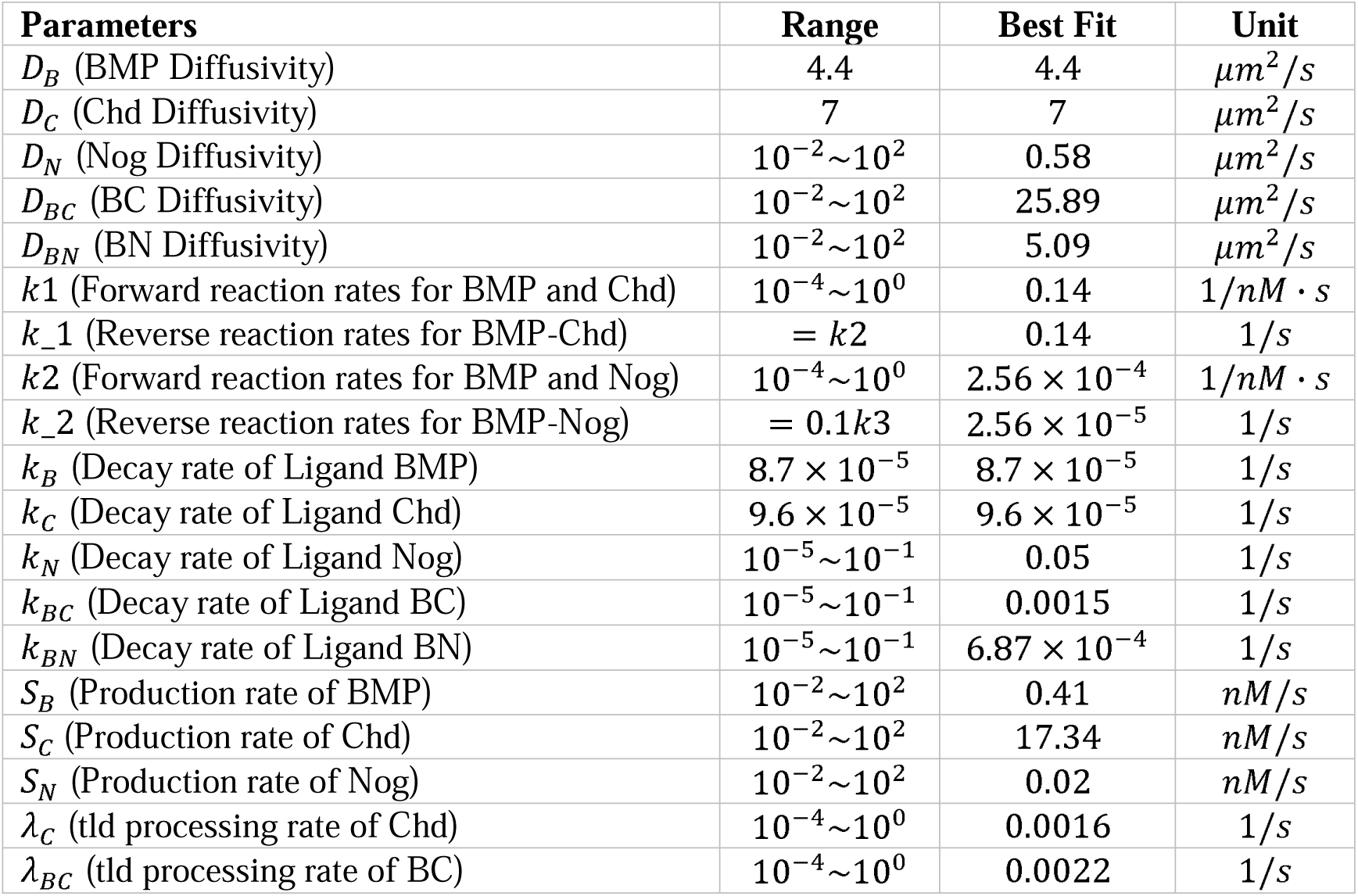
Parameters range.

## Results

We identified the simulations generating BMP profiles that fit the P-Smad5 gradient at 4.7, 5.3, and 5.7 hpf as measured by a low normalized root mean squared deviation (NRMSD) for both WT and Chd LOF. Figure 2A demonstrates a simulation result for a WT case at these time points. The simulated BMP concentration level and measured P-Smad5 profiles (Figure 2B) are normalized between 0 and 1 to calculate the relative error (Figure 2C) between each profile for the entire domain. In our current simulations, we find good agreement throughout the embryo, except on the ventral-anterior side and the lateral region between ventral and dorsal, that exhibits a relative error of ∼12% (Figure 2C). The current best-fit parameter set is listed in Table 2. Compared with the biophysical requirements and fitness from the 1D model screen of Zinski et al., the result is consistent with the source-sink mechanism in which BMP diffuses from its distributed source to a sink of dorsal Chd. Over all, the simulation results match the trend with the experimental P-Smad5 profiles throughout the entire 3D domain. As the simulation processes, the difference accumulates in the ventral-anterior region and at the lateral portion of the margin for later stages. The likely source for this error is the assumption of binary expression domains (Figure 1E-G).

To clarify the differences between simulation results and P-Smad profiles, we isolated the profiles along the embryo’s margin and central region for WT and Chd LOF in Figure 3 for the current best-fit model. The central region is defined as the region connecting over the animal pole the ventral- and dorsal-most points of the embryo. Thus, for different growth stages, the number of indices measured radially along the margin remains the same, but the central region covers increasing radial portions with epiboly. The relative level of BMP ligands on the margin region agree with the experimental P-Smad5 profile for different time points (Figure 3A, 3C). However, the central profile shows a gap for the Chd LOF model. Compared to the experimental profile, which has a nearly linear drop from the ventral to dorsal region, the simulation results’ profiles show a stronger sigmoidal shape. Again, this may be caused by the sharp boundary of BMP production in the lateral region between ventral and dorsal or the sudden appearance of inhibiting Chd ligands. Based on the current data (Zinski et al. 2017) the P-Smad5 gradient in *chordin* mutants showed a statistically significant increase in lateral regions of the embryo during these time points. Our results demonstrate the WT model has relatively better fitness on the central line compared to the Chd LOF model (Figure 3 A-D) where the Chd LOF simulation overestimates at the early time points. The fitness at 4.7, 5.3, and 5.7 hpf for WT exhibits an overall consistency and solutions lay within one standard deviation of the measured mean, however, the earlier 4.7 hpf timepoints between the model predictions and P-Smad data for the Chordin mutants show greater differences along the line that travels through the animal pole (central, Figure 3D).

**Figure 3.**
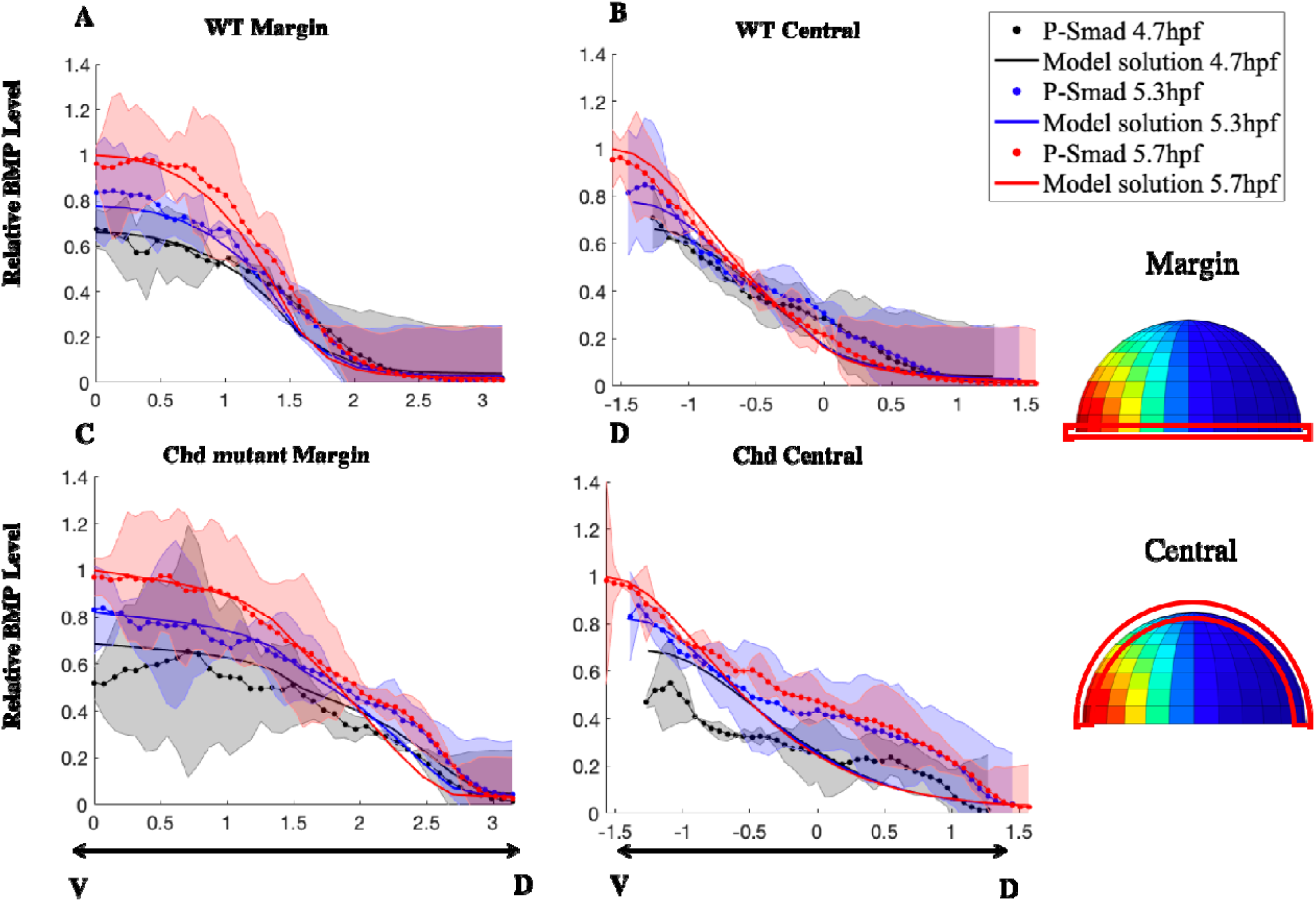
BMP distribution of a single modeling simulation result compared with the P-Smad 5 profile. A, WT case profile in the margin region. B, WT case profile in the central region. C, Chordin LOF case on margin region. D, Chordin LOF case on the central region. All graphs are from ventral to dorsal as indicated below x-axis in C, D.. Black color represents 4.7 hpf, Blue color represents 5.3 hpf, Red color represents 5.7 hpf. The shaded region indicates standard deviation for individual experimental data, represented by the dotted line. The x-axis indicates the radial position measured every degree.

To determine how sensitive the solutions are to the model parameters and identify the likely contributors to the data-model mismatch, we developed a local sensitivity analysis and we also carried out principal component analysis (PCA). In our work, PCA did not identify clear correlations amongst parameters to principal components in models that fit the available data. We relied more heavily then, on parametric sensitivity, and Figure 4 A-D shows the parameter space for different unknown parameters vs. error between normalized simulation results and P-Smad5 averaged intensities. The best parameters among 300,000 random sets were chosen as the starting point to test the local sensitivity. Compared to other unknown parameters, the production-related parameters demonstrate the greatest sensitivity for the simulation error. As shown in Figure 4A, the model fitness changed most in response to BMP and Chd production rate changes, where the smallest error of 5% occurred with BMP production at and Chd at. Interestingly, model fitness does not vary with Nog production rate. The current best fit parameter set is a Chd dominant model, where the production rate of Chd () is much higher than the Nog production rate (). Thus, the WT model sensitivity shows that Chd dominates the sharp BMP gradient profile rather than Nog, consistent with previous 1D modeling results (Zinski et al. 2017).

**Figure 4.**
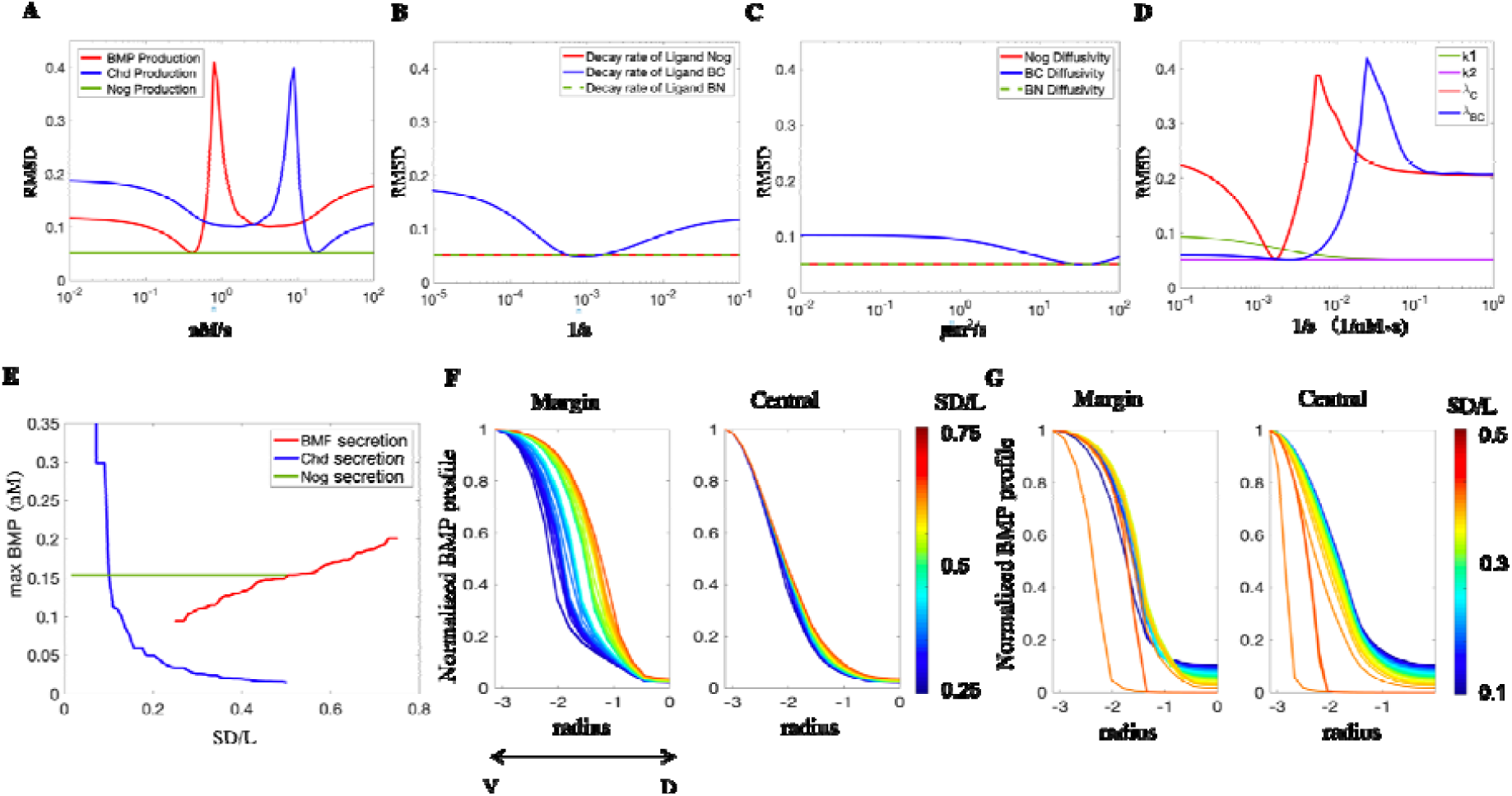
Local sensitivity analysis for unknown parameters. A, production rate for BMP, Chd, and Nog. B, Decay rate for Nog, BMP-Chd complex, and BMP-Nog complex. C, Diffusivity of Nog, BMP-Chd complex, and BMP-Nog complex. D, Tld processing rate of Chd and BMP-Chd complex (1/s), 2^nd^ order constant for BMP/Chd (k1) and BMP/Nog (k2) (1/nM s). For Figure A-D, the x-axis represents the parameters’ value in log scale. The y-axis represents the error of the simulation result for the normalized BMP ligand concentration compared to the normalized experimental P-Smad intensity. E, Maximum BMP concentration changes as secretion domains change with BMP, Chd and Nog. X-axis represents the proportion of secretion domain vs embryo length (SD/L). F. Normalized BMP profile at margin and central line changed by proportional range of BMP secretion. G. Normalized BMP profile on margin and central line changed by proportional range of Chd secretion. X-axis represents the radial position from Ventral (left) to Dorsal (right). Red line indicates a wider range of BMP or Chd secretion, respectively.

Since the decay rates and diffusivities of BMP and Chd are fixed based on measurements (Pomreinke et al. 2017; Zinski et al. 2017), the local sensitivity for the decay rates and diffusivities of Nog, BMP-Chd complex, and BMP-Nog complex are compared in Figure 4B and 4C. The model shows limited sensitivity to the decay rate of BMP-Chd. The model fitness is not sensitive to the decay rate and diffusivity of Nog. The Tld processing rate of Chd and the BMP-Chd complex show similar and relatively large sensitivities. The best Tld processing rate of Chd appears at 1.6 × 10^−3^/*s*, and model fitness worsens considerably when the Tld processing rate of BMP-Chd complex increases beyond 4.1 × 10^−3^/*s*. Also, the model shows less sensitivity to the association constant for BMP/Chd and is insensitive to the association constant for BMP/Nog. The parameters related to Noggin show minimal effects on model fitness based on our current results. On the contrary, the model is sensitive to the Chd-related activities (Chd production, Decay rate of BC complex, BC diffusivity, and Tld processing rate of Chd and BC complex). This also indicates that a precise production region of the major components (BMP, Chordin, Noggin and Tld) and production rate of them could be the critical factors that increases the fitness quality of our current model. The high sensitivity to the current Tld model (linearly dependent on rate and concentration) also suggests that a more detailed protease model is needed to know how Chordin is restricted to the dorsal side required for shaping the BMP gradient.

Since, we use the range of mRNA expression to represent the spatial organization of protein secretion of different components and due to our previous work showing sensitivity to the spatial distributions in the 1D model, we carried out a deeper sensitivity analysis of these terms. To relate the model to the observed phenotypes, we tested how the secretion range of BMP, Chd, and Nog will influence the maximum concentration level of BMP ligands by using the same best-fit parameter set. Shown in Figure 4E, as the range of BMP secretion increases, the maximum BMP ligand concentration increases. On the other hand, as Chd decreases, the maximum concentration level of BMP ligand increases and leads to a dramatic increase when Chd range is less than 10% of the embryo length. Shown in Figure 4F, we compared the changes of the normalized BMP profile in the margin and central region with respect to changes in the ranges of secretion. As BMP expression widens, the BMP ligand concentration profile widens in the margin but does not lead to a substantial difference in the central region. Alternatively, as Chd expression widens, it sharpens the BMP ligand concentration profile and limits the BMP ligand range for both the margin and central region (Figure 4G).

## Discussion

One advantage of simulating the model on the 3D geometry proposed here is that it enables observation of the variation of the level throughout a more realistic whole embryo domain. Though morphogen activity along the margin has offered preliminary insight into DV axis patterning, differences in model fitness are observed along the margin vs. the meridian line that passes through the animal pole.

The major challenge for this study is the vast unknown parameter space and computationally expensive simulation. The 3D growing embryo scheme is considerably more complex compared to a fixed 1D or 3D model. Using the finite difference scheme, the model growth increments are at a constant radial rate along the longitudinal direction and also along a regular shape, which limits our capability to introduce a more realistic embryo shape and incorporate the advection caused by the cell mobility during later epiboly. However, by controlling the mesh size and growth duration in 3D that considers both the spatial and temporal aspects of embryo-scale modeling, this system allows for a closer surrogate to the actual embryo geometry to test different mechanisms that 1D and fixed domain models cannot support.

Local sensitivity analysis provides evidence that the BMP gradient is sensitive to the parameters related to Chordin and Tld activities. Since the BMP gradient is most sensitive to the gene expression inputs for BMP ligand, Chordin, and Tld processing, future work is needed to determine the relative expression of each of these proteins and the relative levels and spatial distributions of their expression. We are collecting *bmp, chordin, noggin*, and *tld* quantitative mRNA wholemount expression profiles at different stages to directly address this question. This work contributes to our long-term goal of developing 3D models of the embryo with growth and advective cell movement, quantitative gene expression, and feedback to determine the interplay of these processes on pattern formation. The study here, is the first step and provides a 3D mathematical model on a growing domain and provides a computational framework to elucidate how the components work together to establish the BMP gradient at multiple time-points in the blastula embryo.

## Supplementary files

MATLAB code for 3D growing domain model and post analysis https://github.com/linlinli12/Zebrafish_BMP_3D_FiniteDifference

## Acknowledgements

We would like to acknowledge funding from NIH grants R01GM132501 to DMU, and NIH R01GM056326 and R35GM131908 to MCM, and editing help from Matt Thompson.

